# Intraspecific chemodiversity provides plant individual- and neighbourhood-mediated associational resistance towards aphids

**DOI:** 10.1101/2022.12.21.521353

**Authors:** Dominik Ziaja, Caroline Müller

## Abstract

1. Some plant species express an extraordinarily high intraspecific diversity in phytochemicals (= chemodiversity). As discussed for biodiversity, higher chemodiversity may provide better protection against environmental stress, including herbivory. However, little is known about whether the resistance of a plant individual towards herbivores is mostly governed by its own chemodiversity or by associational resistance provided by conspecific neighbours.
2. To investigate the role of chemodiversity in plant-aphid interactions, we used the Asteraceae *Tanacetum vulgare*, whose individuals differ pronouncedly in the composition of leaf terpenoids, forming distinct chemotypes. Plants were set-up in a field consisting of 60 plots, each containing five individuals of either the same or different chemotypes. Presence of winged aphids, indicating aphid attraction, and abundance of winged and unwinged aphids, indicating fitness, were scored weekly on each plant, focusing on three commonly occurring aphid species specialised on *T. vulgare*. During the peak abundance of aphids, leaf samples were taken from all plants for re-analyses of the terpenoid composition and quantification of terpenoid chemodiversity, calculated on an individual plant (Shannon index, Hs_ind_) and plot level (Hs_plot_).
3. Aphid attraction was neither influenced by chemotype nor plot-type. The real-time odour environment may be very complex in this setting, impeding clear preferences. In contrast, the abundance was affected by both chemotype and plot-type. On average, more *Uroleucon tanaceti* aphids were found on plants of two of the chemotypes growing in homogenous compared to heterogenous plots, supporting the associational resistance hypothesis. For *Macrosiphoniella tanacetaria* the probability of presence on a plant differed between plot-types on one chemotype. Terpenoid chemodiversity expressed as a gradient revealed negative Hs_plot_ effects on *U. tanaceti*, but a positive correlation of Hs_ind_ with the abundance of *M. tanacetaria*. Aphids of *M. fuscoviride* were not affected by any level of chemodiversity.
4. *Synthesis*. This study shows that not only the chemotype and chemodiversity of individual plants but also that of conspecific neighbours influence plant-herbivore interactions. These effects are highly specific with regard to the plant chemotype, the aphid species as well as its morphs (winged vs. unwinged). Furthermore, our results highlight the importance of analysing chemodiversity at different levels.

## 1 INTRODUCTION

Plants show a huge variation in natural products both interspecifically and intraspecifically, which affects interactions with other organisms across different trophic levels (Schneider et al., 2021, Moore et al., 2014, Glassmire et al., 2016). Such phytochemical diversity, also called chemodiversity (Müller and Junker, 2022), is recently increasingly studied, combining omics approaches with ecological concepts and applying diversity indices developed for biodiversity research (Hilker, 2014, Wetzel and Whitehead, 2020). Plants are confronted with numerous generalist and specialist antagonists, and are thus subjected to a multitude of selective pressures. Across-species comparisons within plant clades revealed that generalist herbivores may promote the synthesis of repellent rather than attractive compounds (Salazar et al., 2018). With more specialised plant-herbivore interactions and at local scales with tight associations the chemical dissimilarity within the plant community tends to increase (Becerra, 2007). Generally, the associational resistance hypothesis postulates that plants in communities with high biodiversity experience lower herbivore infestations than plants in less diverse communities (Randlkofer et al., 2010, Tahvanainen and Root, 1972). The chemodiversity among and within plant species may be a main driver for these effects. Indeed, increasing chemodiversity was found to reduce herbivore damage (Richards et al., 2015, Bustos-Segura et al., 2017), but the outcome also depends on the specific environmental conditions and characteristics of the local herbivore communities (Fernandez-Conradi et al., 2021). In certain plant communities, chemodiversity of low-volatile compounds reduced herbivory by generalists, while specialist herbivory was reduced by high-volatile compounds (Salazar et al., 2016), such as monoterpenoids. So far, research has mostly focused on either community-wide interspecific chemodiversity or individual-based intraspecific chemodiversity, but little is known on the effects of community-wide intraspecific chemodiversity on plant-herbivore interactions (but see, e.g., Bustos-Segura et al., 2017).

Terpenoids are the most diverse group of plant natural products, consist of isoprene (C_5_H_8_) precursors and are often stored in glandular trichomes on the leaf surface of, for example, Asteraceae (Aschenbrenner et al., 2013, Gershenzon and Dudareva, 2007). Terpenoids are either stored and emitted constitutively or induced upon exposure to abiotic or biotic stresses (Gershenzon and Dudareva, 2007). Against herbivores, they can act directly as repellent or deterrent or impact the development and reproduction. However, these compounds can also attract specialists or be involved in indirect plant defence by attracting natural enemies of herbivores (Boncan et al., 2020, Gershenzon and Dudareva, 2007). Plant fitness-enhancing effects of terpenoids may also occur on a community level. Semi-volatile sesquiterpenoids of different hydrophobicity emitted from one plant were found to be adsorbed and re-emitted by its allospecific neighbours, repelling herbivores and thus providing associational resistance, while for a highly volatile monoterpene this effect could not be observed (Himanen et al., 2010). Such mixed blends across neighbouring plants may disturb the host localisation of insects, because they likely use the specific ratio of compounds in volatile blends for this purpose (Bruce et al., 2005). The real-time odour environment of an individual may in the end determine whether it is attracted to its host or distracted by repellent effects (Shao et al., 2021).

In aphids, host localisation is mainly performed by the winged morphs. Compared to unwinged morphs, winged morphs of *Rhopalosiphum padi* were shown to have a higher expression of genes involved in olfactory systems and thus a higher sensitivity towards volatile compounds (Peng et al., 2020), playing a critical role in host colonisation. A peak of winged aphid morphs can usually be observed in early summer when colonisation takes place in a more dispersed pattern. Subsequently, unwinged aphids become more dominant and colonisation occurs more locally (Mehrparvar et al., 2013, Dixon, 1977, Braendle et al., 2006). Despite winged morphs being potentially more sensitive to volatile plant metabolites, both morphs can be affected by metabolites located on the leaf surface, in the leaf and in the phloem sap (Wink et al., 1982, Chang et al., 2022, Powell et al., 2006). Terpenoids have been shown to attract or repel aphid species (Kos et al., 2013, Webster et al., 2008), while terpenoids stored in glandular trichomes negatively affected fecundity and longevity of aphids (Wang et al., 2021). Apart from the terpenoids, the nutritional value of the phloem sap is particularly important for aphids and determines whether aphids will stay on the plant and be able to reproduce (Karley et al., 2002, Nowak and Komor, 2010).

The perennial Asteraceae common tansy, *Tanacetum vulgare* L., is an aromatic plant species that shows a pronounced intraspecific variability of mono- and sesquiterpenoids (Judzentiene and Mockute, 2005, Keskitalo et al., 2001, Wolf et al., 2011), which are passed on genetically (Holopainen et al., 1986, Keskitalo et al., 2001). Based on their relative composition, terpenoids are used to assign tansy individuals to specific mono-chemotypes with one predominant terpenoid or mixed-chemotypes with one to two additional major satellite terpenoids (Holopainen et al., 1986). In nature, different chemotypes often occur together in patches next to each other (Kleine and Müller, 2011). Tansy is visited by numerous herbivores (Schmitz, 1998), including various aphid species that are mono- or oligophagous on tansy. The chemotypes have been revealed to play a crucial role in interactions between tansy and its specialised herbivores (Kleine and Müller, 2011, Benedek et al., 2019, Jakobs and Müller, 2018). For example, winged morphs of *Metopeurum fuscoviride* preferably colonise chemotypes emitting α-thujone, (*E*)-dihydrocarbone, α-copaene and β-cubebene (Clancy et al., 2016), whereas unwinged morphs of *Uroleucon tanaceti* use most likely terpenoids to locate preferred plant parts within a plant individual (Jakobs and Müller, 2019). Moreover, tansy chemotypes also affect aphid populations directly by impacting their growth rate (bottom-up) and indirectly by influencing the establishment of different arthropod food webs (top-down), resulting in chemotype-specific communities of aphids, their predators and mutualistic ants (Senft et al., 2018, Balint et al., 2016). In turn, aphids can also modulate the nutritional quality of the phloem sap with changes depending on the aphid species, the plant part as well as the chemotype (Jakobs et al., 2019). Thus, this plant species offers a highly suitable model to study the role of chemodiversity in plant-aphid interactions.

This study investigated how certain leaf chemotypes and varying degrees of intraspecific chemodiversity on both individual plant and group level (= direct neighbourhood) of tansy plants affect the attraction and occurrence (presence and abundance on a host plant) of aphids specialised on this plant species. A common garden experiment was set up containing homogenous and heterogenous groups (= plots) of five plant individuals per plot belonging to one or five chemotypes and aphid presence per plant was scored across the season. Chemodiversity was considered on the plant individual level and on the plot level. We hypothesised, that (1) certain chemotypes are more attractive to winged aphids than others and (2) that homogenous chemotype plots are more attractive than heterogenous plots, potentially due to a less mixed odour bouquet facilitating host localisation in the former and associational resistance in the latter. Moreover, we expected that (3) chemotype and plot-type in interaction affect attraction towards and presence and abundance on the host plants differently depending on the aphid species, and (4) plants and plots with a lower chemodiversity enhance the occurrence of aphids on plant individuals, used as a proxy for aphid fitness.

## 2 MATERIALS AND METHODS

### 2.1 Experimental set-up

A stock of tansy plants was established from seeds collected in January 2019 at four different sites in Bielefeld, Germany (51°58’58.52’N, 8°27’12.27’E; 51°58’51.8’N, 8°27’40.0’E; 51°58’59.3’N, 8°28’13.8’E; 51°58’42.2’N 8°28’35.5’E; elevation 98-105 m). Seeds were collected from 8-10 mother plants per site with at least 20 m distance between each plant, to enhance the likelihood to collect seeds from genetically distinct individuals (tansy can grow clonally). Seeds were germinated and terpenoid profiles determined from young leaves as described in 2.3. Based on the monoterpenoid profiles, plants of five distinct chemotypes were picked to build up the stock collection. The terpenoid composition of two of these chemotypes was dominated (> 55% of total terpenoid concentration) by a single monoterpenoid (mono-chemotypes), either artemisia ketone (called “Keto” chemotype in the following) or β-thujone (“BThu”). The composition of the other three chemotypes was dominated (10 – 50% of total terpenoid concentration) by two to three compounds (mixed-chemotypes), either α-thujone and β-thujone (“ABThu”), artemisyl acetate, artemisia ketone and artemisia alcohol (“Aacet”) or (*Z*)- myroxide, santolina triene and artemisyl acetate (“Myrox”) (in total *n* = 30 plants per chemotype, originating from different mother plants, called chemo-genotypes hereafter). Two clones from each of these plants were produced from rhizome cuttings, grown in a greenhouse and transferred outside into a mixed soil:sand-bed from September to December 2019 for acclimatisation, and kept there until the last week of May 2020. Then, these non-flowering plants were introduced into a field common garden.

The common garden was set up close to Bielefeld University (52°03’39.43’N, 8°49’46.66’E; elevation 142 m). Of each of in total 150 chemo-genotypes, one clone was planted in a homogenous plot consisting of five plants of the identical chemotype (*n* = 30 plots; 6 plots per chemotype), the other in a heterogenous plot (*n* = 30 plots) consisting of one plant of each of the five chemotypes. The site (24 x 17 m) was split into six blocks, each consisting of ten plots (1 x 1 m) with 1 m between plots and 2 m between blocks (for detailed set-up see Supplement Fig. S1). Plants were planted in PVC-tubes (diameter 16 cm, height 30 cm), which were inserted 25 cm deep into the soil to allow for distinction of individual plants (referred to as pots from here on). Within plots, pots were arranged in a circle with 72° between neighbouring plants. All plants within a plot were descendants from different maternal plants.

The area between blocks and plots was milled once in spring and once in autumn every year. Throughout the season, vegetation within plots was removed manually if it reached half the length of the maximum tansy plant height in the respective plot. When occurring, seedlings of additional tansy plants were removed. In December 2020, the aboveground biomass of all plants was harvested down to 5 cm above ground, thereby homogenising plant growth once and potentially removing overwintering aphid eggs, thus enhancing the chance that plants were colonised by migrating aphids in 2021.

### 2.2 Scoring of aphids

Scoring of aphids visiting the experimental tansy plants took place on a weekly basis from May 6^th^ 2021 until August 18^th^ 2021, when almost no aphids were observed anymore. It was performed usually midweek within two days between 8 am and 6 pm. For scoring of aphid species specialised on tansy, each plant was carefully examined, trying to prevent the aphids from dropping off the plant. Winged and unwinged individuals per species were counted separately. The presence (yes/no) of winged aphids on a plant or plot was considered as “attraction”. Observing at least one aphid of any morph on a plant individual was considered as species presence (yes/no). Aphid counts (=abundance) were taken as an indication of aphid fitness. Because *U. tanaceti* develops huge colonies, extra measures were implemented: (a) counting was capped when a colony on a plant reached 2000 individuals; (b) starting on June 30^th^ aphid populations with a count ≥ 100 were estimated in increments of ten. To account for the variance explained by the presence of ants for ant-tended aphid species (*M. fuscoviride*), the presence of ants in plant pots and ants actively harvesting honeydew from aphids were noted down. Furthermore, ant nests in the ground either within or directly around the plant pot were scored after manually checking for ant activity by poking into the nest using a stick. Every week, the sampling order and person counting (*n* = 5) were randomised on the plot-level.

### 2.3 Plant phenotyping

On June 21^th^-22^th^, all plants were phenotyped, assessing the length of the highest shoot, the number of shoots, the number of leaves and the number of stems with inflorescences per plant. For later analyses of the actual terpenoid composition (see 2.4), the top 4 cm of the youngest fully developed, non-infested leaf of one of the stems was cut off and immediately frozen in liquid nitrogen. Leaf samples were harvested between 10 am and 1:30 pm. Most except ten plants had not developed flowerheads at that time point. One plant of a heterogenous plot turned out to belong to a different chemotype than originally suggested. Thus, all data for this entire plot was excluded from all statistical analyses.

### 2.4 GC-MS analysis of leaf terpenoid composition

The harvested leaf material was freeze-dried, homogenised, weighed and extracted in heptane containing 1-bromodecane (Sigma Aldrich, Taufkirchen, Germany) as an internal standard in an ultrasonic bath for 5 min. Samples were centrifuged and the extracts analysed using gas chromatography coupled with mass spectrometry (GC-MS; GC 2010plus – MS QP2020, Shimadzu, Kyoto, Japan) on a semi-polar column (VF-5 MS, 30 m length, 0.2 mm ID, 10 m guard column, Varian, Lake Forest, United States) in electron impact ionisation mode at 70 eV and with helium as carrier gas. Samples were injected at 240 °C with a 1:10 split. A starting temperature of 50 °C was kept for 5 min, ramping up to 250 °C at 10 °C min^-1^, then increasing with 30 °C min^-1^ to a final temperature of 280 °C, hold for 3 min. Blanks of heptane with the internal standard and an alkane standard mix (C7–C40, Sigma Aldrich, Taufkirchen, Germany) were measured regularly between sample batches. Terpenoids were identified based on their Kovàts retention indices (KI) (Kovàts, 1958) and by comparing spectra to synthetic reference compounds, where available, and to entries of the libraries NIST (National Institute of Standards and Technology, Gathersburg, USA, 2014), Pherobase (El-Sayed, 2012) and those reported in Adams (2007). Terpenoids were quantified based on the peak area of the total ion chromatogram and the relative composition determined by dividing each peak area by the sum of the peak areas of all peaks within each sample.

### 2.5 Statistical analyses

All statistical analyses were carried out in R version 4.2.1 (R Core Team, 2022), using the packages vegan (Oksanen et al., 2022), dplyr (Wickham et al., 2021), glmmTMB (Brooks et al., 2017), DHARMa (Hartig, 2021), car (Fox and Weisberg, 2019), insight (Lüdecke et al., 2019), emmeans (Lenth, 2021), pgirmess (Giraudoux, 2022) and glmnet (Friedman et al., 2010). Visualisations were made in ggplot2 (Wickham, 2016). A virtual environment of the RStudio project is available on github (https://github.com/DoZi93/CommonGarden_DataAnalysis).

The occurrence of winged aphids (attraction) per plant and total count (abundance) per week were used for further analyses. To exclude zero-values simply due to the absence of an aphid species, all data recorded for every species (winged presence, total count) was filtered across weeks based on the cumulative sum over the season; the first week exhibiting ≥ 1% was selected as first, the week displaying the elbow of the curve towards the end of the season was selected as last week. If nymphs of *U. tanaceti* with signs of wing-forming (see 2.2) were observed on a plant during a counting event, their count was included in the total aphid count but set to zero for all analysis targeting winged aphid count and presence, because they were likely produced by aphids that had already colonised the plant. For all generalised linear mixed models (GLMM) of total counts, goodness of fit and the appropriate distribution (Poisson, negative binomial 1, negative binomial 2) were evaluated based on simulated residuals using DHARMa plots. Occurrence of winged aphids was binary data and therefore analysed using binomial distribution. Random effects causing convergence problems due to low variance explained were dropped from the models.

Non-transformed whole season total count data of every species was analysed using zero-inflated generalised linear mixed models (zi-GLMM) after checking for zero-inflation. The aphid presence (at least one winged or unwinged individual observed on plant) was modelled by the zero-inflation component of the total count models. In all models, chemotype, plot-type, the chemotype x plot-type interaction and calendar week were implemented as fixed effects. Since *M. fuscoviride* is ant-tended, presence of ants, ant nests and ants actively tending aphids were included as binary, fixed effects for the models related to this aphid species. Block, plot number, plant clone ID, plant ID, maternal genotype and observer were included as random effects in every model. Nestedness of random effects was accounted for by uniquely coding the nested random effects (block and plot number). The zero-inflation model component was modelled with the same fixed and random effects as the conditional model.

In addition, for the week in which plant morphological and chemical data were sampled, effects of plant traits on winged aphid presence and the total count of aphids were analysed on the individual plant- and plot-level using non-zero inflated GLMMs. The individual plant-level chemodiversity Hs_ind_ was quantified by calculating the Shannon diversity index *H_S_* = -Σ *p*_i_ * ln *p*_i_, (Shannon and Weaver, 1964), with *p* being the relative abundance of each terpenoid within an individual. For an individual plant-level analysis, ln (x+1)-transformed morphological parameters, namely length of highest shoot, number of stems, number of leaves and number of stems with inflorescences, as well as Hs_ind_ were included in the models as fixed effects and the same random effects, except for plant ID, were used as in the whole season GLMMs. To assess effects on the plot-level, data was summarised for each plot: presence of winged aphids was evaluated plot-wise. Total aphid count, number of leaves, number of shoots and number of shoots with inflorescences were summed up per plot; for the length of the highest shoots the average across plants per plot was taken. The plot chemodiversity Hs_plot_ was calculated with *p* being the average of the relative abundance of each terpenoid across the five plants per plot. Except for the aphid-related response variables and the Hs_plot_, variables calculated on plot-level were also ln (x+1)-transformed. The fixed effects were identical to those used in the individual plant-level models, while only block was included as random effect.

#### 2.5.1 Importance of single terpenoids on aphid occurrence

To infer the effects of individual terpenoids on aphid-tansy interactions, a Poisson-LASSO regression was applied to those total aphid counts, which were significantly affected by the chemodiversity (*Macrosiphoniella tanacetaria* by Hs_ind_, *U. tanaceti* by Hs_plot_), using the relative composition of each terpenoid from the individual plant (individual plant-level) or the average of each plot (plot-level) as predictor variables (Salazar et al., 2018). The penalisation term integrated in a LASSO regression allows for coefficients to be estimated zero, resulting in both feature selection and assessment of correlations between features and response variables (James et al., 2013). The winged aphid presence was analysed using a binomial LASSO regression, the total count of aphid species using a Poisson LASSO regression. Minimal lambda of all LASSO models was determined using K-fold cross validation.

## 3 RESULTS

### 3.1 Effects of chemotype and plot-type on aphid attraction and occurrence

Across the season, the tansy specialist aphid species *U. tanacetaria*, *M. tanaceti* and *M. fuscoviride* were frequently found on the experimental plants. The attraction of winged aphids and the presence and total count of winged and unwinged morphs of these aphid species depended on the calendar week (*X^2^* ranging from 9.03 to 57049, *p* < 0.001) with the exceptions of winged morph presence and total presence of *M. fuscoviride* (Table 1). On average, higher numbers of *U. tanaceti* were observed on homogenous plots of the Keto and ABThu chemotype, whereas on the Myrox chemotype numbers were higher when this chemotype grew in heterogenous plots (Figs. 1a, 2a, Table 1). In contrast, not the total count but the presence of winged and unwinged morphs of *M. tanacetaria* was affected by the chemotype x plot-type interaction (Fig. 1b, Table 1), i.e. there was a higher probability to observe the aphid species on ABThu plants in homogenous compared to heterogenous plots (Fig. 2b). The total count of *M. fuscoviride* was positively affected by the presence of ants tending their colonies (model predictions: 3.70 ± 1.21 without ant-tending, 18.94 ± 6.14 with ant-tending), but not by plot-type nor chemotype (Fig. 1c, Table 1). Furthermore, across all whole season models the mother plants explained less than 1% of the variance of the datasets (Table 1).

**Figure 1:**
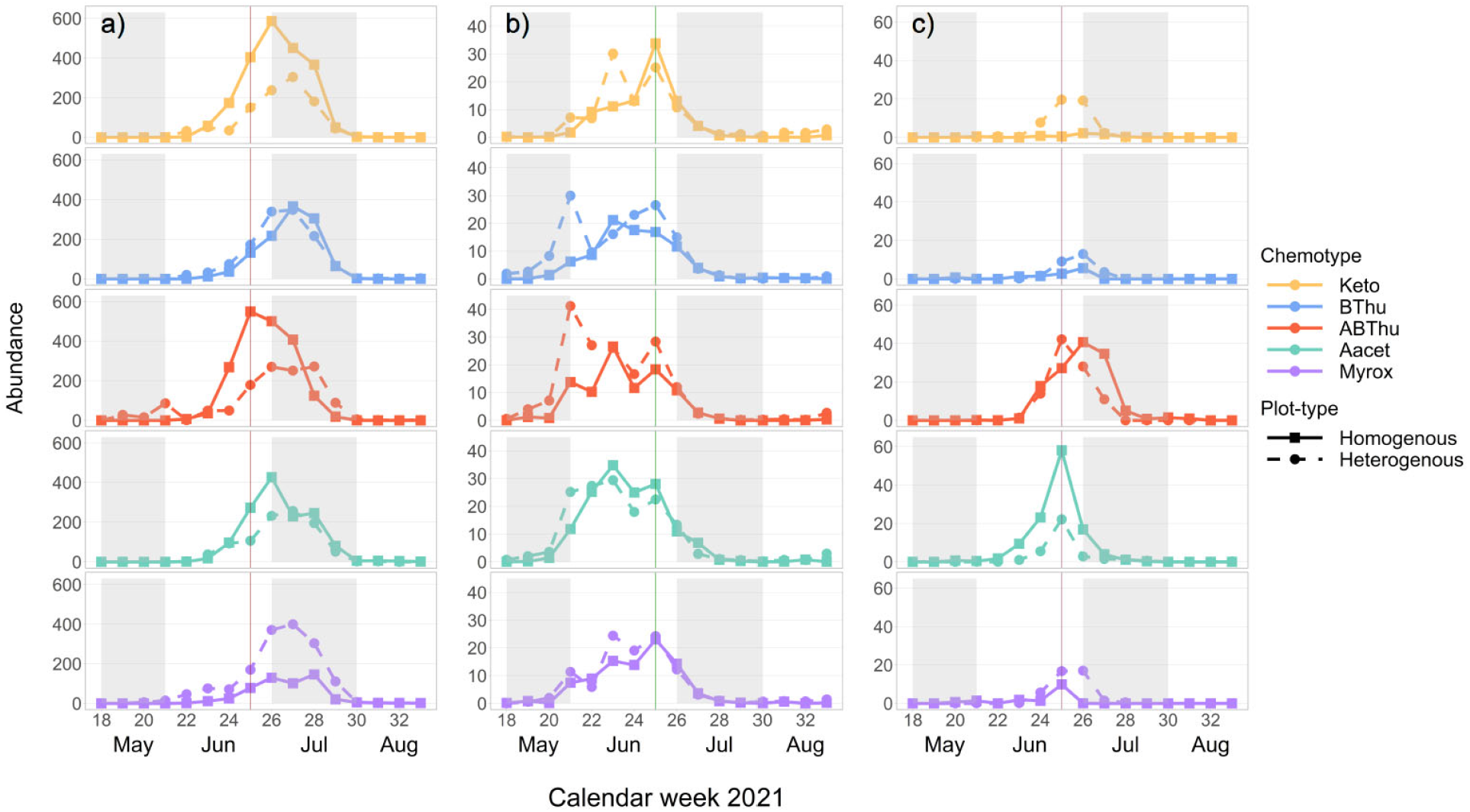
Mean total count of winged and unwinged aphids of a) *Uroleucon tanaceti*, b) *Macrosiphoniella tanacetaria* and c) *Metopeurom fuscoviride* per calendar week on plants of *Tanacetum vulgare* grown in plots of five individuals of the same (homogenous) or different (heterogenous) chemotypes (Keto: artemisia ketone chemotype, BThu: β-thujone chemotype, ABThu: α-/β-thujone chemotype, Aacet: artemisyl acetate/artemisia ketone/artemisia alcohol chemotype, Myrox: (*Z*)- myroxide/santolina triene/artemisyl acetate chemotype) across the season. Vertical lines indicate the calendar week in which the morphological and leaf terpenoid characterisation of the plants took place; *n* = 25-30 per chemotype x plot-type combination.

**Figure 2:**
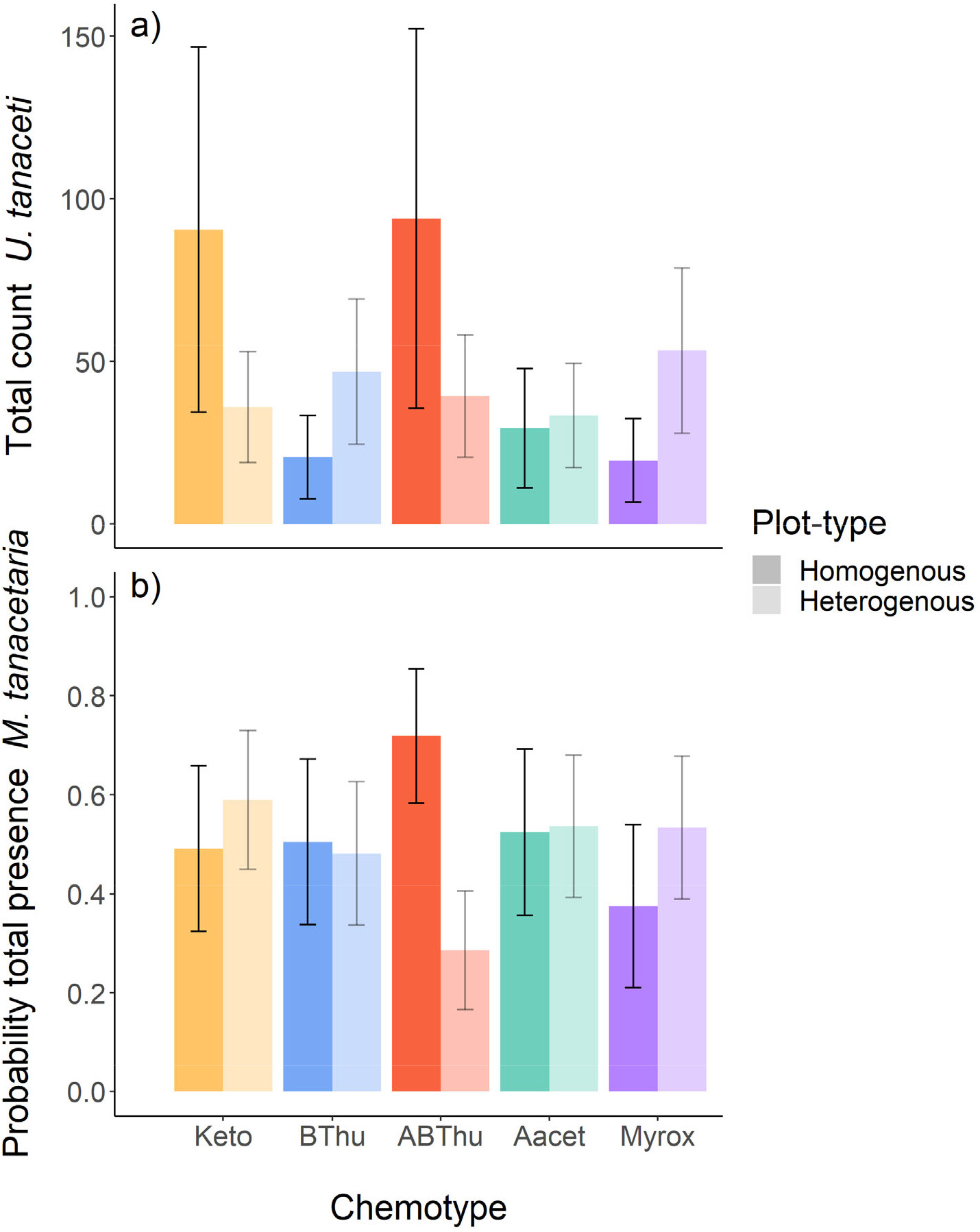
Em mean predictions of (a) the total count of winged and unwinged individuals of *Uroleucon tanaceti* and (b) the probability of presence of winged and unwinged individuals of *Macrosiphoniella tanacetaria* on plants of *Tanacetum vulgare* grown in plots of five individuals of the same (homogenous) or different (heterogenous) chemotypes (Keto: artemisia ketone chemotype, BThu: β-thujone chemotype, ABThu: α-/β-thujone chemotype, Aacet:artemisyl acetate/artemisia ketone/artemisia alcohol chemotype, Myrox: (*Z*)-myroxide/santolina triene/artemisyl acetate chemotype). A significant (a: *p* = 0.01, b: *p* = 0.02) chemotype x plot-type interaction was found based on the conditional (a) and zero-inflation (b) component of the zi-GLMM. Number of observations per chemotype x plot-type interaction *n*_obs_ = 200-240 and plants per chemotype x plot-type interaction *n*_plants_ = 25-30.

**Table 1:**
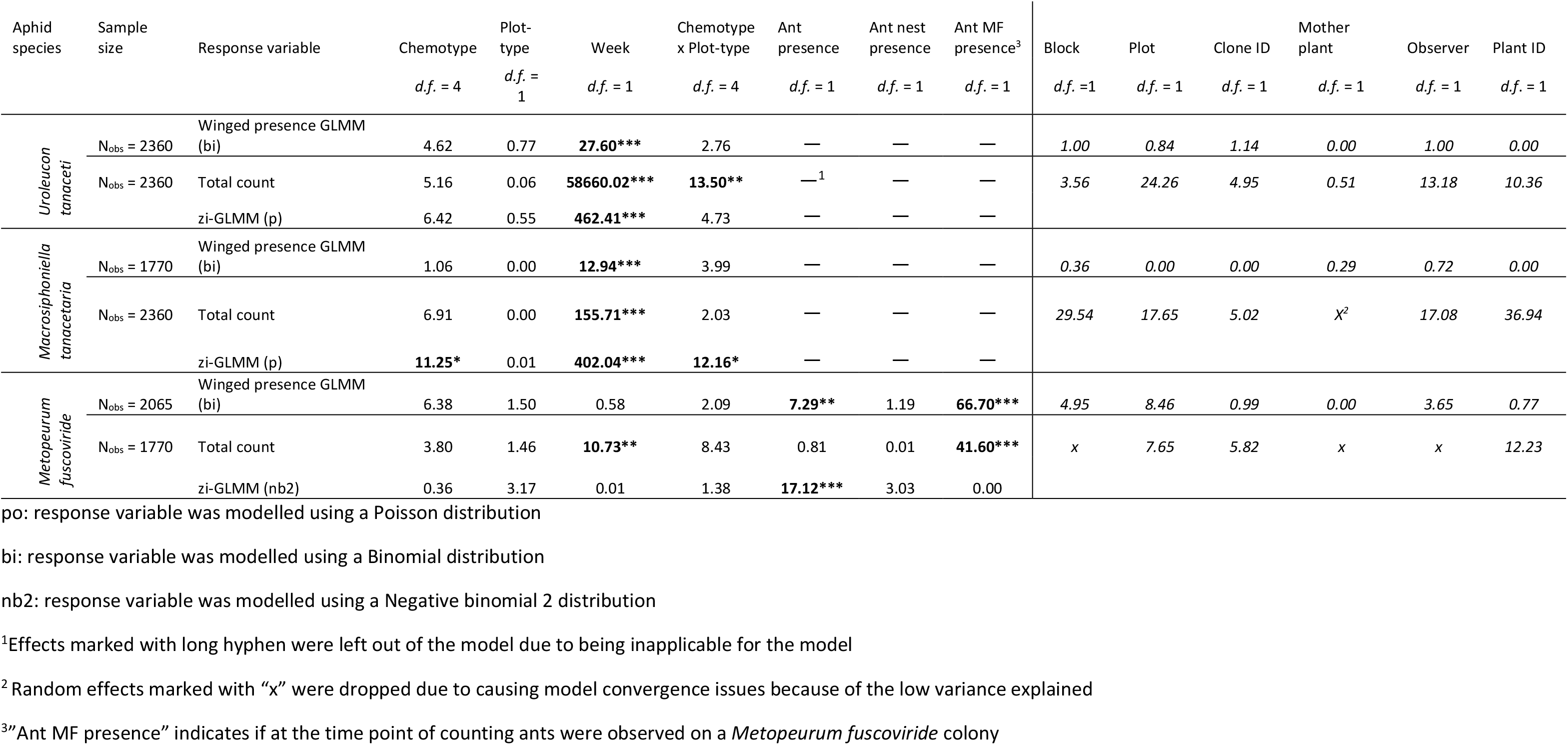
(zi-)GLMM estimates of total count (unwinged and winged aphids) and presence of winged morphs of three aphid species specialised on *Tanacetum vulgare* across the counting season. Shown are the *X*^2^ estimates based on Wald’s type 3 chi-square test (regular font) and the proportion of variance explained by random effects (italic). Numbers in bold font indicate significant effects with asterisks displaying the significance level (* *p* < 0.05; ** *p* < 0.01; *** *p* < 0.001). For zero-inflation models estimates of the conditional model are displayed in the first line, estimates of the zero-inflation model in the second line.

### 3.2 Terpenoid composition and chemodiversity Hs_ind_ and Hs_plot_

In total, 52 terpenoids were found in leaves throughout all plants (Table S1). The monoterpenoid(s) originally dominating each chemotype at initial terpenoid analyses continued to represent > 50% of the terpenoid composition per chemotype (Fig. 3). Some monoterpenoids dominating in one chemotype could also be found across multiple chemotypes. For example, the Keto chemotype showed on average small proportions of *α*-thujone (6.5%), while the Aacet chemotype contained santolina triene (10.1%) as well as (*Z*)-myroxide (16.0%). Other monoterpenoids present in higher proportions were sabinene (BThu 7.9%, ABThu 4.5%) and 1,8-cineole (ABThu 4.4%, Santo 3.5%, BThu 3.1%). Regarding sesquiterpenoids, in every chemotype γ-cadinene (8.3 - 10.8%) and an unknown sesquiterpenoid (KI 1673, 1.0 – 4.9%) were present.

**Figure 3:**
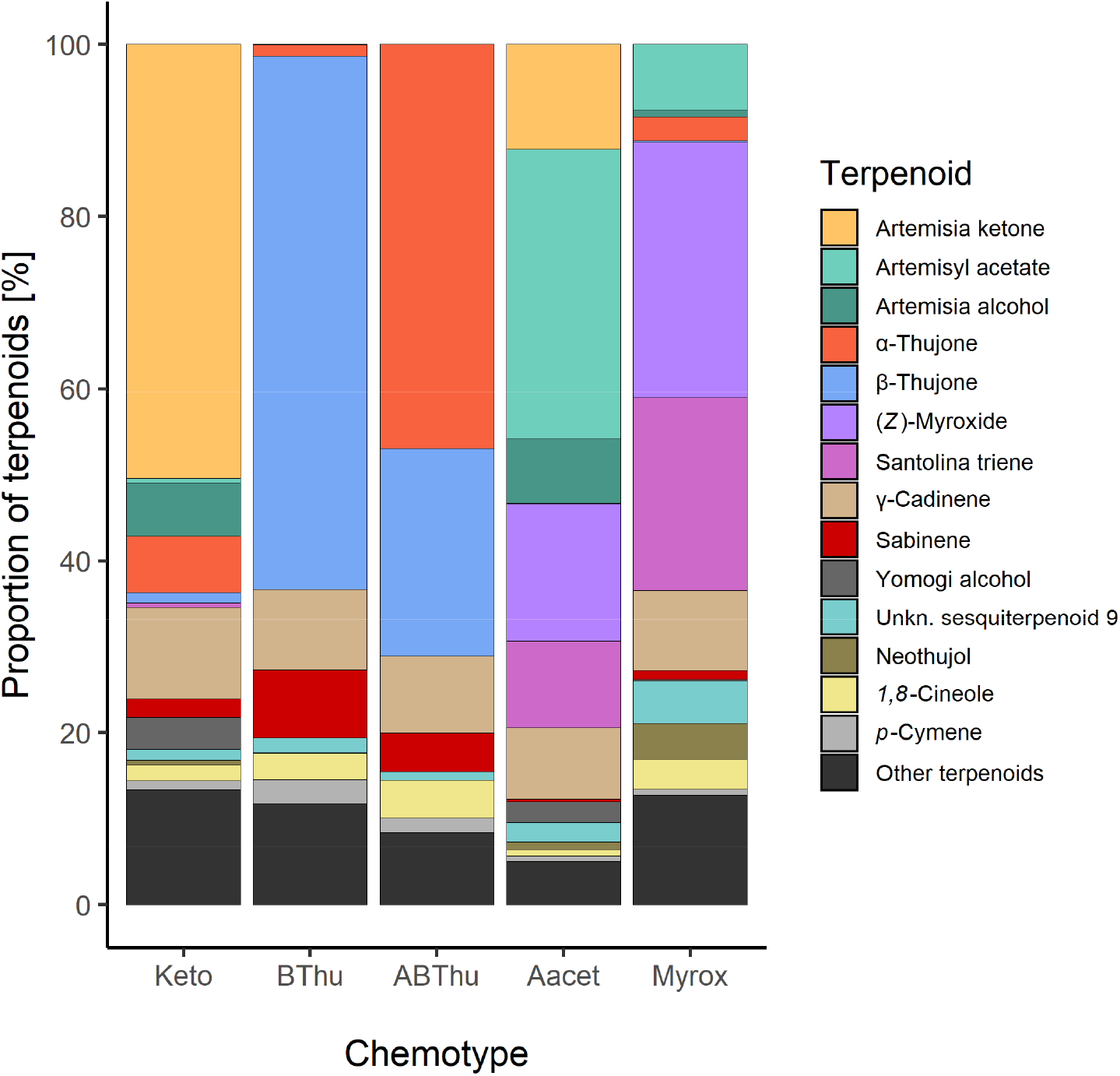
Mean terpenoid composition of leaves of *Tanacetum vulgare* of five different chemotypes grown in the common garden, sampled in June 2021. All terpenoids are individually displayed that account together for ≥ 80% of the total chemotype composition; *n* = 55-60 individuals per chemotype.

In homogenous and heterogenous plots the average Hs_ind_ ranged between 1.16 and 2.01, with the lowest value found in plants of the BThu chemotype grown in heterogenous plots. On the plot level, the average Hs_plot_ was with 2.51 highest, when plants were grown in heterogenous plots (Table 2).

**Table 2:**
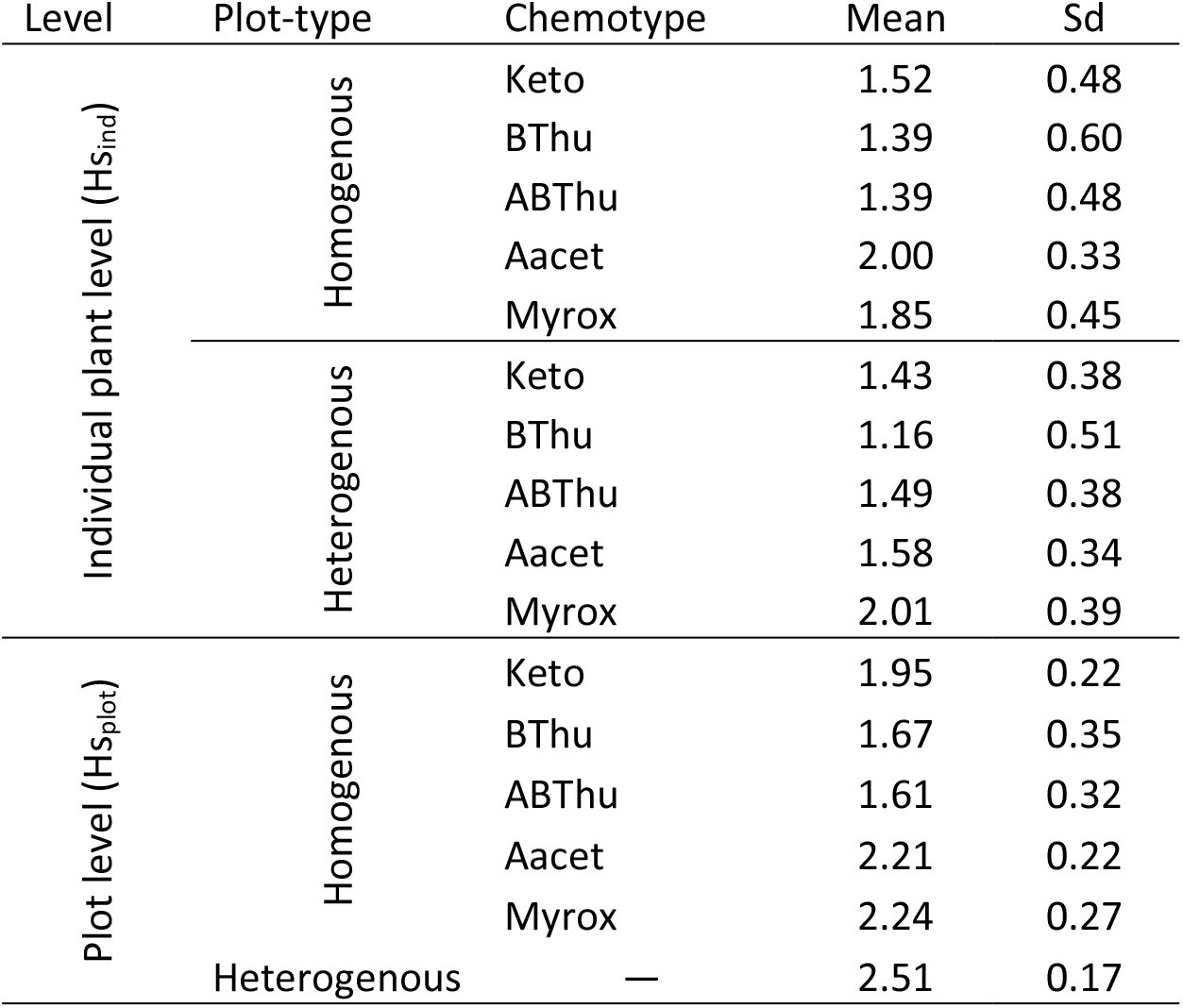
Shannon-diversity (Hs) of the five tested chemotypes calculated based on the relative terpenoid composition obtained from GC-MS measurements of youngest, fully developed leaf samples taken in calendar week 25 from each plant planted in the common garden setup. Mean and standard deviation (Sd) are calculated across either each individual plant (Hs_ind_; *n* = 25-30 per chemotype x plot-type combination) or the average across five plants per plot (Hs_plot_; *n* = 4-5 per homogenous chemotype and heterogenous plots). For composition of chemotypes see Fig. 3.

| Level | Plot-type | Chemotype | Mean | Sd |
| --- | --- | --- | --- | --- |
| Individual plant level ($H_{s_{ind}}$) | Homogenous | Keto | 1.52 | 0.48 |
|  |  | BThu | 1.39 | 0.60 |
|  |  | ABThu | 1.39 | 0.48 |
|  |  | Aacet | 2.00 | 0.33 |
|  |  | Myrox | 1.85 | 0.45 |
|  | Heterogenous | Keto | 1.43 | 0.38 |
|  |  | BThu | 1.16 | 0.51 |
|  |  | ABThu | 1.49 | 0.38 |
|  |  | Aacet | 1.58 | 0.34 |
|  |  | Myrox | 2.01 | 0.39 |
| Plot level ($H_{s_{plot}}$) | Homogenous | Keto | 1.95 | 0.22 |
|  |  | BThu | 1.67 | 0.35 |
|  |  | ABThu | 1.61 | 0.32 |
|  |  | Aacet | 2.21 | 0.22 |
|  |  | Myrox | 2.24 | 0.27 |
|  | Heterogenous | — | 2.51 | 0.17 |

### 3.3 Effects of individual plant- and plot-level morphology as well as Hs_ind_ and Hs_plot_ on aphid occurrence and attraction

In the week in which leaf sampling took place, winged aphids of *U. tanaceti*, *M. tanacetaria* and *M. fuscoviride* were observed on 162, 82 and 28 of the 291 plants from which terpenoids were re-analysed, respectively. On the individual plant-level, the likelihood of observing winged aphids (attraction) of *U. tanaceti* was reduced by the number of shoots (*X*^2^ = 3.99; *p* = 0.046) but increased by the number of leaves (*X*^2^ = 7.29; *p* = 0.01). Winged aphids of *M. tanacetaria* and *M. fuscoviride* were not affected by any morphological plant trait on both individual plant- and plot-level (Table S2). Moreover, Hs_ind_ and Hs_plot_ did not show any effect on the presence of winged aphids (Table S2).

The total count (abundance) of both *U. tanaceti* and *M. fuscoviride* was positively affected by the number of leaves on the individual plant-level. Regarding the plot-level, plant shoots were more influential than the leaves: *U. tanaceti* was positively affected by the shoot height, *M. fuscoviride* was negatively affected by the number of shoots, while numbers of leaves had no effect. Next to morphological traits, the total count was also affected by plant chemodiversity in aphid species-specific ways. On the individual level, Hs_ind_ affected the total count of *M. tanacetaria* positively (Table 3, Fig. 4a). In contrast, on the plot level Hs_plot_ had a negative influence on the total aphid count of *U. tanaceti* (Table 3, Fig. 4b). For both of these significant effects, the LASSO regression estimated the coefficients of all 52 individual terpenoids to be zero, i.e., no linear combination of terpenoids could predict these two counts.

**Figure 4:**
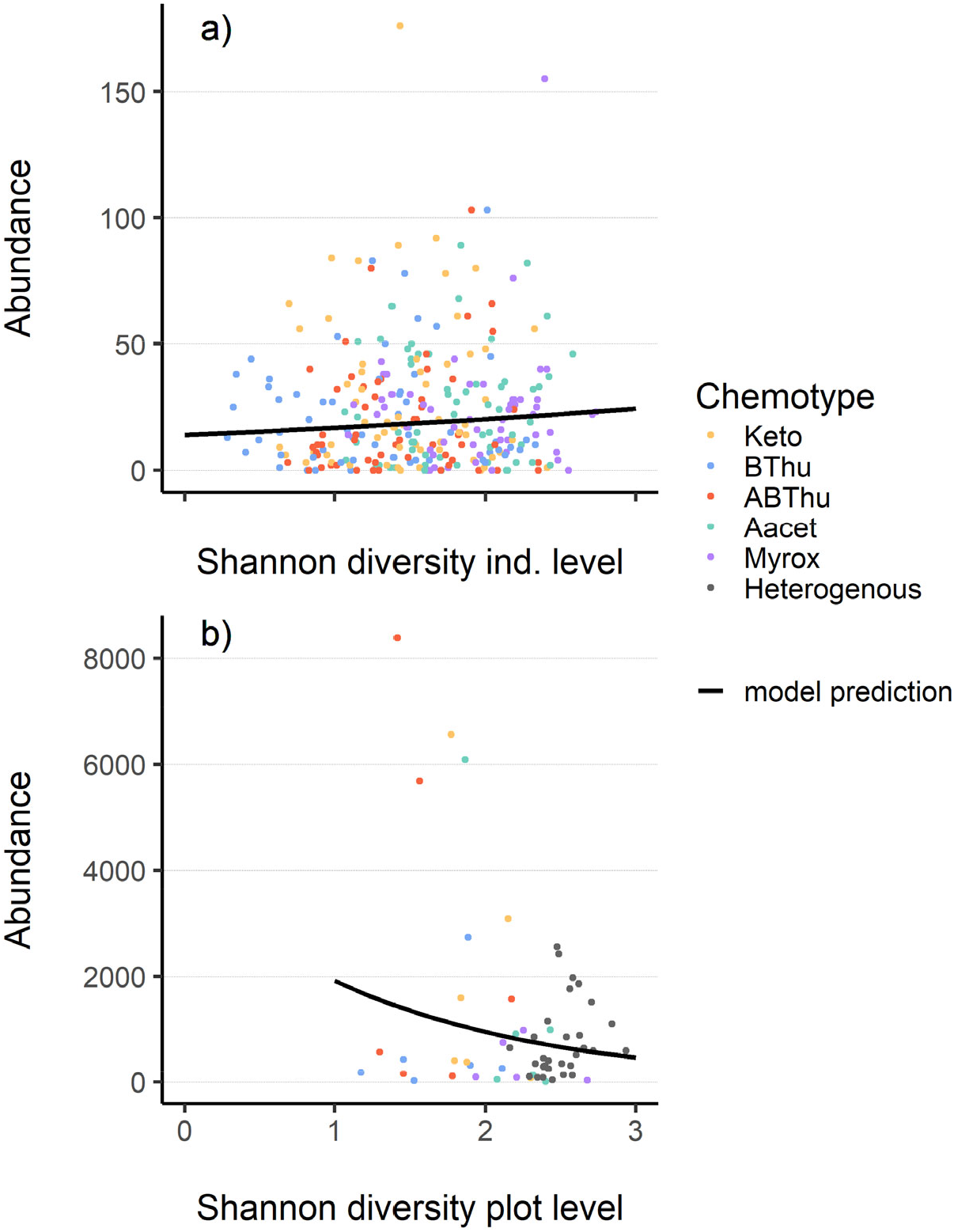
Relationship between Shannon diversity of terpenoids in leaves on a) plant individual level (Hs_ind_) or b) plot level (Hs_plot_) and total count of aphids (winged and unwinged) in June 2021 on *Tanacetum vulgare* plants of five different chemotypes grown in plots of five individuals of the same (homogenous) or different (heterogenous) chemotypes (Keto: artemisia ketone chemotype, BThu: β-thujone chemotype, ABThu: α-/β-thujone chemotype, Aacet: artemisyl acetate/artemisia ketone/artemisia alcohol chemotype, Myrox: (*Z*)-myroxide/santolina triene/artemisyl acetate chemotype). (a) Count of *Macrosiphoniella tanacetaria* on every individual plant; *n*plants = 55-60 per chemotype. (b) Cumulative count of *Uroleucon tanaceti* for every plot; *n*plots = 5-6 (per homogenous plots) or 30 (heterogenous plots).

**Table 3:** GLMM-estimates of total count (winged + unwinged morphs) of aphid species specialised on *Tanacetum vulgare* in calendar week 25 of 2021. The X^2^ estimates are based on Wald’s type 3 chi-square test, numbers in bold font indicate significant effects, asterisks display the significance level (* p < 0.05; ** p < 0.01; *** p < 0.001). Results of winged presence were moved into the supplement since no significant effects were found. Total number of plants on which aphids were scored: 291.

| Response variable |  | *Uroleucon tanacetii* |  |  |  | *Macrosiphoniella tanacetaria* |  |  |  | *Metopeurum fuscoviride* |  |  |  |
| --- | --- | --- | --- | --- | --- | --- | --- | --- | --- | --- | --- | --- | --- |
|  |  | Total count (abundance)<sup>b</sup> |  |  |  | Total count (abundance)<sup>a</sup> |  |  |  | Total count (abundance)<sup>a</sup> |  |  |  |
| Level |  | Individual plant level |  | Plot level |  | Individual plant level |  | Plot level |  | Individual plant level |  | Plot level |  |
| **Fixed Effects** | *d.f.* | $X^2$ | Effect | $X^2$ | Effect | $X^2$ | Effect | $X^2$ | Effect | $X^2$ | Effect | $X^2$ | Effect |
| Shannon diversity | 1 | 0.61 | - | **3.90\*** | ↓<sup>5</sup> | **3.90\*** | ↑ | 0.15 | - | 1.00 | - | 0.25 | - |
| Shoot length | 1 | 0.87 | - | **8.28\*\*** | ↑ | 0.02 | - | 3.17 | - | 2.08 | - | 0.25 | - |
| Number of shoots | 1 | 0.03 | - | 0.00 | - | 0.02 | - | 0.004 | - | 0.01 | - | **10.29\*\*** | ↓ |
| Number of leaves | 1 | **16.24\*\*\*** | ↑<sup>4</sup> | 0.31 | - | 3.31 | - | 0.45 | - | **4.75\*** | ↑ | 0.94 | - |
| Number of shoots with inflorescences | 1 | —<sup>2</sup> | — | — | — | 0.14 | - | 0.00 | - | 0.15 | - | 1.94 | - |
| Ant presence | 1 | — | — | — | — | — | — | — | — | **52.49\*\*\*** | ↑ | **14.57\*\*\*** | ↑ |
| **Random Effects** |  | Variance [%]<sup>3</sup> |  | Variance [%] |  | Variance [%] |  | Variance [%] |  | Variance [%] |  | Variance [%] |  |
| Block | 1 | 2.72 |  | 17.65 |  | 18.65 |  | 21.50 |  | 0.00 |  | 0.00 |  |
| Plot | 1 | 29.65 |  | — |  | 13.77 |  | — |  | 21.24 |  | — |  |
| Mother plant | 1 | 0.00 | | — | | 1.24 | | — | | $X^6$ | | — | |

<sup>a</sup>response variable was modelled using a negative binomial 1 distribution
<sup>b</sup>response variable was modelled using a negative binomial 2 distribution
<sup>1</sup> $p$ -values were estimated using a type-3 Wald chi-square test
<sup>2</sup>Effects marked with long hyphen were left out of the model due to being inapplicable for the model
<sup>3</sup>Variance [%] of random factors displays the percentage of the total variance that is explained by the random factors
<sup>4</sup>↑ display a positive relationship with the fixed effect, <sup>5</sup>↓ displays a negative relationship with the fixed effect
<sup>6</sup>Random effects marked with “x” were left out of the model due to causing model convergence problems

## 4 DISCUSSION

We assessed the effects of intraspecific chemodiversity of plants, here in the form of differently composed leaf terpenoids, on the presence of winged aphids, indicating attraction, as well as total aphid count, indicating fitness. These effects were assessed on the individual plant as well as on the plant neighbourhood level. Overall, our results showed that chemodiversity mainly affected the presence and abundance of aphids on the plants, but not the attraction of winged aphids. Furthermore, these effects were highly specific both in terms of the plant chemotype and the aphid species.

The attraction of all three detected aphid species specialised on tansy was neither affected by the chemotype, plot-type, nor their interaction, in contrast to our expectations. First, we hypothesised that aphids would find certain chemotypes more attractive than others. Evidence for clear preferences for certain chemotypes by *U. tanaceti* and *M. tanacetaria* comes from laboratory assays, in which unwinged aphids were offered choices between leaves of two chemotypes in different combinations (Jakobs and Müller, 2019, Jakobs and Müller, 2018). Aphid choice seems more clear-cut in such restricted conditions than in the field, where several other differences between plant individuals and numerous other environmental factors make the real-time odour environment highly complex, particularly for flying morphs. Nevertheless, higher early colonisation by winged morphs of another aphid species was found in natural conditions for chemotypes emitting α-thujone among other terpenoids (Clancy et al., 2016). Winged morphs of another aphid species feeding on Poaceae displayed higher gene expression levels of chemosensory proteins than unwinged ones (Peng et al., 2020), suggesting that high sensitivity towards odour profiles may be expected particularly in these morphs. Second, we expected a higher attraction of aphids to plants in homogenous plots since these should emit less-mixed odours than heterogenous plots. Once the different plots are approached by the winged individuals, cues deciphering host occupancy, providing information about prior aphid (Mehrparvar et al., 2014) or virus infestations (Eigenbrode et al., 2002), or visual cues (Döring and Chittka, 2007) might become more important for attraction to — or repellence of — certain plant individuals than the volatile terpenoid cues. Such diverse factors may explain, why no preferences for chemotype or plot-type were observed in the present study. Apart from such effects, the distance between plants of different chemical profiles may also be decisive for distinct preference behaviour of winged aphids, as the mixing degree of different plant odours will be affected (Shao et al., 2021). Recently it has been revealed that the exact ratio of compounds in the odour bouquet of a plant is already changing at short distances (Cai et al., 2022), which may affect host plant discrimination, particularly in stands of conspecific neighbourhoods.

No effect on the pure presence but lower counts, i.e. abundances, of winged and unwinged *U. tanaceti* were observed on plants of the Keto and ABThu chemotype in homogenous compared to heterogenous plots, while the opposite pattern was found on plants of the Myrox chemotype. These results suggest that plants of the latter chemotype are more resistant towards *U. tanaceti* than the other tested chemotypes. Also under laboratory conditions, aphids of *U. tanaceti* showed different population growth on plants of different chemotypes (in interaction with plant part) (Jakobs and Müller, 2018), indicating that chemical differences between these plants can be important factors influencing aphid fitness. Apart from host-plant related factors, also top-down effects may have influenced the aphid counts. Indeed, tansy is known to establish chemotype-specific arthropod food webs (Balint et al., 2016). Our results clearly show that not only the individual plant chemotype but also the chemotype composition in the neighbourhood of the respective plant is of importance in plant-herbivore interactions. The lower abundance of aphids on plants of the Keto and ABThu chemotype in heterogenous compared to homogenous plants supports the associational resistance hypothesis (Randlkofer et al., 2010, Tahvanainen and Root, 1972), which predicts lower herbivore damage in more diverse communities. Diverse neighbourhoods may also harbour more natural enemies, leading to reduced herbivore pressure (Cappuccino et al., 1998). Such associational effects apparently also apply on plant communities of conspecifics that are chemically diverse and may explain why we find high intraspecific chemodiversity in some species. Surprisingly, the associational resistance seems specific for certain chemotypes. For instance, the BThu and Aacet chemotype showed no differences in the abundance of *U. tanaceti* between the plot-types, whereas abundances were even higher on plants growing in heterogenous plots for the Myrox chemotype. Neighbouring tansy plants may benefit in a highly chemotype-specific way from top-down effects, as predators also show different abundances on distinct chemotypes of tansy (Benedek et al., 2019). Moreover, in our specific set-up all chemotypes may benefit from and/or provide associational resistance towards other antagonists than *U. tanaceti*. Another possible mechanism for associational effects is the enrichment of leaves from neighbouring plants with compounds of lower volatility such as sesquiterpenoids (Himanen et al., 2010). The sesquiterpenoids α-humulene and β-caryophyllene, of which the latter was present in higher proportions in the well-defended Myrox tansy chemotype, were found to reduce longevity and fecundity of aphids in tomato (Wang et al., 2021). Whether tansy indeed also adsorbs volatiles from neighbours, which may affect herbivores, remains to be tested.

Comparing our findings between the three aphid species, our hypothesis that the effects of chemotype in interaction with plot-type on the presence and abundance are species-specific is supported. In contrast to *U. tanaceti*, no effects were observed on the abundance of *M. tancetaria*, but the probability of presence of winged and unwinged aphids was higher on plants of the ABThu chemotype in homogenous compared to heterogenous plots, supporting again associational resistance in this chemotype. When growing in homogenous plots, plants of this chemotype may be particularly supportive for *M. tancetaria*. A higher presence of aphids may lead to an enhanced release of volatiles due to induction, causing even higher colonisation of these plants for aphids, but this idea needs further testing. Impacts of intraspecific differences in groups of metabolites on arthropods and different inducibility have been found for other plant metabolite classes in other plant species. For example, differences in glucosinolate profiles of different populations of *Brassica oleracea* influenced the performance of both herbivores and their parasitoids (Harvey et al., 2011), and those plants showed specific pattern in herbivore-induced concentrations of glucosinolates (Harvey et al., 2011, Gols et al., 2018). Moreover, for aphids, different morphs can also show differences in both physiology and behaviour, even towards the same volatiles (Webster, 2012, Peng et al., 2020). Unwinged morphs may favour a broader range of host cues and are attracted to more volatiles than winged morphs, potentially due to the fact that they do not migrate and thus encounter less non-host cues (Webster, 2012). In our experiment, unwinged aphids may have been most likely offspring from winged aphids that had settled and reproduced but we cannot exclude that also some unwinged aphids moved between plants standing close to each other. In particular, *M. tanacetaria* readily drop from their host plants when being disturbed (personal observation) and may then switch to a close-by neighbour. In contrast to the two other aphid specialists on tansy, the presence and abundance of *M. fuscoviride* was not affected by chemotype or plot-type, but total counts were higher in the presence of ants tending aphid colonies. This finding is in line with previous findings that report positive effects of ant-tendance on the reproduction and other life history traits of *M. fuscoviride* (Flatt & Weisser, 2000). In another study significant effects of tansy chemotypes dominated by either *L*-camphor, (*Z*)-β-terpineol or eucalyptol on the abundance of *M. fuscoviride* were found (Senft et al., 2018). Thus, tritrophic interactions between plants, aphids and ants may be chemotype-specific. Season affected almost all traits of the three aphid species studied here, with peak abundances differing slightly between species. Seasonal variation in plant chemistry may contribute to this pattern. In addition to this temporal differentiation, the three species also show spatial niche differentiation by colonising different plant parts, with *U. tanaceti* infesting old leaves, *M. tanacetaria* young plant parts (Jakobs et al., 2019) and *M. fuscoviride* prefering stems and inflorescences (Loxdale et al., 2011). Thus, although competition between these three species was found (Mehrparvar et al., 2018), they can co-exist.

Attraction and occurrence of the three aphid species were also analysed with respect to the terpenoid chemodiversity calculated as a gradient rather than distinct chemotype-classes. The sampling for this terpenoid profile analysis took place during the peak occurrence of the three aphid species in June and thus data were exclusively related to the aphid traits in that particular week. Chemodiversity, calculated as Shannon index (Hs), had no effects on the presence of winged morphs; however, overall, for *M. tanacetaria* and *M. fuscoviride* only few colonisation events by winged aphids took place in this week. In contrast, increasing Hs affected the total count of *U. tanaceti* negatively on a plot-level (Hs_plot_), but not on an individual level (Hs_ind_). Increased chemodiversity has been found to be associated with a decreased herbivore damage in several studies (Salazar and Marquis, 2022, Bustos-Segura et al., 2017, Richards et al., 2015). For example, on *Piper* species particularly negative effects for plots with high levels of high-volatility chemodiversity were found on specialist herbivores (Salazar et al., 2016). The fact that negative effects similar to those observed in interspecifically diverse plots were likewise found for Hs_plot_ of conspecific plots in our tansy system, further supports the associational resistance hypothesis in the scope of intraspecific chemical variation.

Interestingly, for *M. tanacetaria* again a different pattern was found. Here, the total count increased with higher Hs_ind_, while Hs_plot_ showed no significant correlation. The increased Hs_ind_ might in this case not be cause but a consequence of an increased total *M. tanacetaria* count, potentially due to herbivore-induced changes in terpenoids. In fact, an aphid-species specific change in chemical composition has been found in tansy of different chemotypes (Jakobs et al., 2019), but effects on terpenoid profiles by aphid infestation remain to be investigated in this system. Since no correlation was found between *M. tanacetaria* abundance and Hs_plot_, infestation with *M. tanacetaria* may not mediate communication between neighbouring plants to induce chemical defences, as found in other species (Baldwin et al., 2006), or the relationship between the infestation and responses is non-linear. Moreover, we only calculated the chemodiversity of terpenoids, but several other metabolites, particularly of the phloem sap, are highly relevant for determining the performance of aphids (Cao et al., 2018, Dreyer and Campbell, 1987). Given the different effects on different aphid species on tansy, the mechanisms how chemodiversity in itself provides associational effects might not only be specific to the herbivore morph (winged vs. unwinged) and species, but also to the different metabolites and metabolite classes the host plant species expresses.

In conclusion, we showed that intraspecific chemodiversity in plant communities can provide associational resistance against herbivores. Besides, even within specialised herbivore species of the same feeding guild, such resistance is highly specific to the species and even its different morphs. To unravel these effects, it is important to analyse each species interaction, considering also developmental stage, with plant chemodiversity in two different forms on two different levels. Firstly, plant chemodiversity can be measured in the form of distinct classes as chemotypes with their specific composition of plant compounds but also in the form of a continuous gradient as a measure of chemodiversity, where different classes of metabolites could be considered. With sufficient knowledge of biosynthetic pathways, more informative indices could also be used (Petrén et al., 2022). Secondly, chemodiversity can be considered on the level of the individual plant but also summarised on the level of the plant community. In this study, the focus was on the effects of chemodiversity on plant antagonists. Future studies will focus on effects of associational resistance on plant performance by studying growth and fitness traits in these chemically more or less diverse neighbourhoods.

## ACKNOWLEDGEMENTS

We thank numerous co-workers of the Chemical Ecology Department being involved in the set-up of the common garden, in particular Lukas Brokate, Stephanie Champion, Elisabeth E. Eilers, Ruth Jakobs and Rohit Sasidharan. Moreover, we thank Liv Krause, Tobias Rützert, Tim Kirrmann and Vanessa Schellenberg for helping to score the aphids and Elisabeth E. Eilers, Liv Krause and Thuan Nguyen for the chemical analysis of the terpenoids. For the preparation and heavy maintenance of the common garden area, we thank Gebhard Sewing. This project was funded by the German Research Foundation (DFG), project MU 1829/28-1, as part of the Research Unit (RU) FOR 3000. We thank all members of the RU for fruitful discussions.

## AUTHORS’ CONTRIBUTIONS

C. M. conceived the ideas and designed the common garden set-up; D. Z. collected and analysed the data and wrote a first draft of the manuscript; C. M. revised the manuscript. All authors gave final approval for publication.

## DATA AVAILABILITY STATEMENT

The raw and processed field data (aphid scores, morphological characterisation, terpenoid analysis) are stored together with the R scripts in the github repository https://github.com/DoZi93/CommonGarden-aphid-2021 in the folder “/Data”. The repository will be made publicly accessible once the manuscript was accepted.

