## Supplement_Fig_S1 for "Intraspecific chemodiversity provides plant individual- and neighbourhood-mediated associational resistance towards aphids"

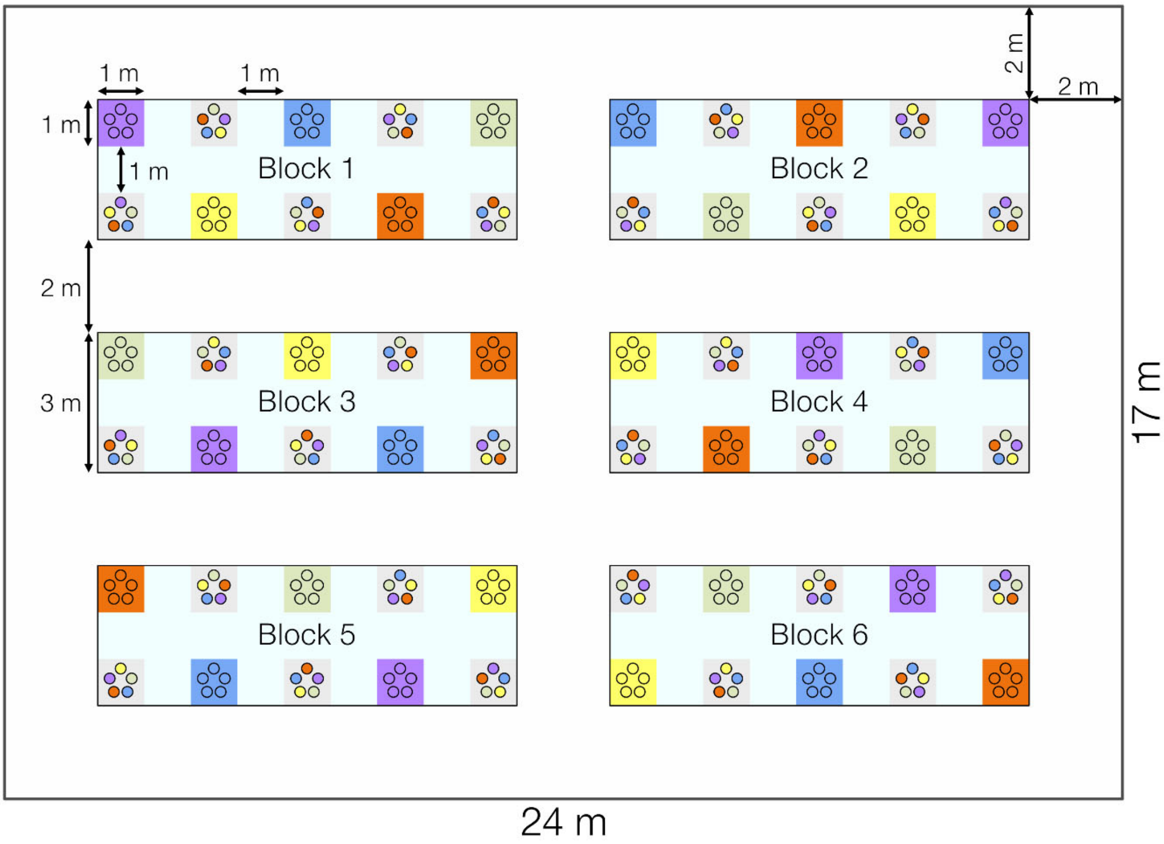


**Figure S1**: Experimental common garden design. Plants of five different chemotypes (yellow – Keto: artemisia ketone chemotype, blue – Bthu: β-thujone chemotype, orange – ABThu: α-/β-thujone chemotype, green – Aacet: artemisyl acetate/artemisia ketone/artemisia alcohol chemotype, purple – Myrox: (*Z*)-myroxide/santolina triene/artemisyl acetate chemotype) were planted in plots with all five individuals sharing the same chemotype (homogenous, unicolored rectangles) or all five individuals expressing a different chemotype (heterogenous, grey rectangles with five differently coloured circles). The experimental set-up was replicated in six blocks. Number of plants in total = 300. As one plant in a homogenous plot turned out to be not the correct chemotype, only 295 plants (59 plots) were considered for further analysis.
